# Citrus-specific truncation of CsGPX4 disrupts intracellular ROS homeostasis

**DOI:** 10.64898/2026.02.13.705763

**Authors:** Shannon Barry, Xin Wang, Satya Swathi Nadakuduti, Nian Wang

## Abstract

Glutathione peroxidases (GPXs) are key antioxidant enzymes that contribute to cellular redox homeostasis through reactive oxygen species (ROS) detoxification and modulation of redox signaling pathways. Although GPX functions have been studied in many annual plants, their roles in woody perennial crops remain poorly understood. Here, we characterized the *GPX* gene family in *Citrus sinensis* and investigated the function of *CsGPX4*, the closest homolog of *Arabidopsis thaliana* AtGPX8 because of its potential role in mediating oxidative stress signaling. Overexpression of CsGPX4 induced an oxidative stress phenotype that includes chlorosis, growth inhibition, cellular degeneration, and elevated intracellular ROS accumulation. Immunoblot and MALDI-TOF analyses showed that *CsGPX4* overexpression resulted in a truncated proteoform, while the full length CsGPX4 was able to accumulate in *Nicotiana benthamiana*, indicating a species-specific post-translational truncation. The truncated CsGPX4 proteoform exhibited an altered subcellular localization shifting from cytoplasmic and nuclear localization to membrane-associated compartment in citrus cells. The truncated form of CsGPX4 had a reduced glutathione peroxidase activity compared to the full-length protein. Proteomic profiling suggested reprogramming of pathways associated with redox detoxification, cytoskeleton organization, hormone signaling, and stress responses. Our study identifies a citrus-specific post-translational regulatory mechanism that modulates CsGPX4 localization and enzymatic activity. This suggests that citrus may employ a species-specific proteolysis processing as a protective mechanism to maintain redox homeostasis during oxidative stress.

## Introduction

Reactive Oxygen Species (ROS), including superoxide anion (O₂•⁻), hydrogen peroxide (H₂O₂), hydroxyl radicals (•OH), and singlet oxygen (¹O₂) are reactive derivatives of molecular oxygen generated during aerobic metabolism (Halliwell and Gutteridge, 2015). ROS are primarily produced in chloroplasts, mitochondria, peroxisomes, and the exocellular side of plasma membrane (Bailey-Serres and Mittler, 2006; Mittler, 2017). While ROS function as key signaling molecules regulating plant growth, development, immunity, and acclimation responses (Mittler et al., 2011; Inzé et al., 2012; Ortega-Galisteo et al., 2012; Baxter et al., 2014; del Rio and Puppo, 2015), their excessive accumulation can damage cellular components including nucleic acids, proteins, lipids, and membranes (Mittler et al., 2022). Therefore, the dynamic maintenance of intracellular redox balance permitting ROS to function in signaling while limiting oxidative damage, known as redox homeostasis, is essential for plant survival and adaptation under adverse conditions (Mittler, 2017; Mittler et al., 2022; Alazem and Burch-Smith, 2024).

Plants have evolved a complex antioxidant defense systems to regulate ROS accumulation through coordinated enzymatic and non-enzymatic mechanisms (Wang et al., 2024). The enzymatic components consist of superoxide dismutase, catalase, ascorbate peroxidase, glutathione peroxidase (GPX), monodehydroascorbate, dehydroascorbate, glutathione S-transferases, glutathione reductase, and alternative oxidases (Wang et al., 2024). Peroxiredoxin and thioredoxin (TRX) also contribute to ROS detoxification through thiol-based redox reactions and antioxidant signaling pathways (Sachdev et al., 2021). The non-enzymatic antioxidant components include ascorbic acid, tripeptide glutathione, phenolics, α-tocopherol, carotenoids, flavonoids, and solute proline that help detoxify ROS by free-radical chain reaction interruption (Sachdev et al., 2021).

GPXs are an important peroxidase that are known to maintain cellular redox homeostasis (Singh et al., 2022). For example, Arabidopsis GPX3 is a an important antioxidant that scavenges excessive H_2_O_2_ while simultaneously severing as an oxidative signal transducer to relay the H_2_O_2_ signal required for abscisic acid and drought stress signaling (Miao et al., 2006). Multiple GPX studies have been carried out in Arabidopsis (Rodriguez Milla et al., 2003). In addition, knockout of *GPX5* in rice (*Oryza sativa*) resulted in salt stress sensitivity, impaired seed germination, and defects in plant development (Wang et al., 2017). In human and animals, GPX catalytic reaction relies on glutathione (GSH) as a reducing agent and contains a selenocysteine residue at the active site (Trenz and Margis-Pinheiro, 2025). In contrast, plant GPXs utilize TRX rather than GSH and possess three conserved non-selenium Cys residues in the active sites (Herbette et al., 2007a; Trenz and Margis-Pinheiro, 2025). Plant GPXs function in three steps involving the oxidation of active-site Cys residue by hydrogen peroxide or organic hydroperoxides, forming a sulfenic acid; the sulfenic acid reacting with a second, nearby cysteine residue to form an intramolecular disulfide bridge; lastly, reducing the disulfide bond by TRX, thus regenerating the active enzyme (Wang et al., 2024).

While antioxidant enzymes are induced in response to elevated ROS levels, their activity and accumulation are regulated through multiple mechanisms, including transcriptional regulation, post-transcriptional control, post-translational modifications (PTMs), retrograde signaling, and protein-protein interactions (Jiménez et al., 2025). Proteins can either undergo reversible or irreversible PTMs and they are known to be tasked with plant metabolite production because they are either enzymes or enzyme regulators (Friso and van Wijk, 2015). PTMs are involved in the changes in oligomeric state, stabilization/degradation, and deactivation (Huber and Hardin, 2004). The irreversible PTM results in the new carboxyl and amino termini formation within cleaved substrates (Fernandez-Fernandez et al., 2023). The proteolysis processes involve the loss, gain or alteration of protein function and play an important role in plant signaling pathway regulations (Cao et al., 2019; Paulus and Van der Hoorn, 2019; Jobin et al., 2020; Liu et al., 2020). Acting as a “molecular switch”, they can either activate or inactivate a specific cellular process (Fernández-Fernández et al., 2023). There have been studies where plant proteases are involved in organellar protein import (Friso and van Wijk, 2015), programmed cell death (Salvesen et al., 2016; Buono et al., 2019), growth and development (Schaller, 2004; Liu et al., 2018), and responses to abiotic and biotic stressors (Balakireva and Zamyatnin, 2018; Salguero-Linares and Coll, 2019). There has been new evidence that suggests that proteolytic processing is an important regulatory mechanism in citrus stress and immune responses. For example, citrus papain-like cysteine proteases (PLCPs) were recently shown to specifically cleave pathogen-derived protein during Huanglongbing (HLB) defense responses (McClelland et al., 2026), highlighting the role of protease-mediated processing in regulating stress-associated signaling pathways.

In our study, we investigated CsGPX4, a *C. sinensis* homolog of Arabidopsis AtGPX8 that has previously shown to be associated with enhanced oxidative stress tolerance whereas loss of function impaired root development and stress resilience (Gaber et al., 2012). CsGPX4 was selected for functional citrus characterization. Despite the importance of GPXs in plant stress responses, the post-translational regulation and functional dynamics of GPX in woody crops remain poorly understood. Unexpectedly, biochemical and proteomic analysis of CsGPXs overexpression lines showed accumulation of a citrus-specific truncated proteoform that altered subcellular localization, impaired enzymatic activity, disrupted intracellular ROS balance, and led to cellular stress phenotypes. These findings suggest that citrus may have a specialized post-translational regulatory mechanism that can modulate protein function under imbalanced systems.

## Results

### Identification and structural characterization of the CsGPX genes family

To identify *GPX* genes in *C. sinensis* and two other citrus species of *C. clementine* and *C. ichangensis*, we conducted a BLAST search using eight GPX proteins from *A. thaliana* as queries. Four *GPX* genes were identified in *C. sinensis* and were designated as *CsGPX1-CsGPX4*, *C. clementine* has five GPX and *C. ichangensis* has four. A phylogenetic analysis based on citrus GPX amino acid sequences resulted in five distinct GPX groups (I-V) based on clustering pattens. (**Fig. 1A**; **Supplementary Table 1**). AtGPX6 did not cluster within any of the defined groups and instead formed a distinct branch, suggesting a possible divergence from other members of the citrus GPX family. Three of the four CsGPX genes (CsGPX1, CsGPX2, and CsGPX4) reside on chromosome 5 whereas CsGPX3 is located on chromosome 4 (**Fig 1B**).

**Figure 1.**
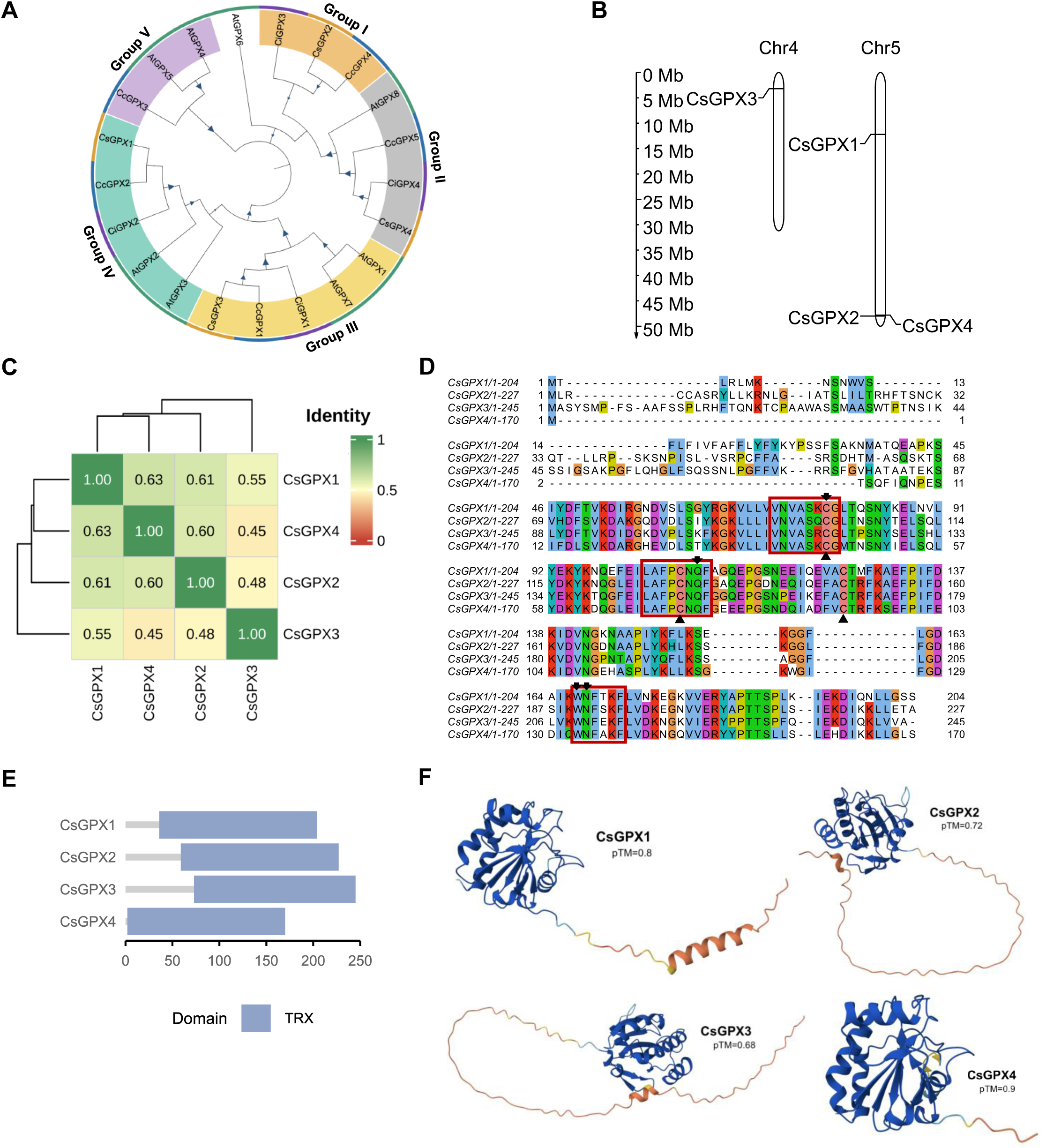
Characterization of the CsGPX proteins in *C. sinensis*. (**A**) Neighbor-Joining phylogenetic tree of *C. sinensis* (orange), *C. clemintime* (blue), *C. ichangensis* (purple) and Arabidopsis (green) proteins, constructed using MEGA-X and iTOL with 1,000 bootstrap replicates. Size of blue triangle represents bootstrap values. Amino acif sequence similarity is represented by groups that is color shaded; group I (orange), group II (grey), group III (yellow), group IV (teal), and group V (purple). No shading represents no group identity. (**B**) Chromosomal positions of *CsGPX1-CsGPX4* in *C. sinensis*. Genomic positions were obtained from the *C. sinensis* reference genome and visualized using custom annotation scripts. (**C**) Pairwise amino acid identity analysis of CsGPX proteins. (**D**) Multiple sequence alignment of CsGPX proteins generated with Clustal omega. The catalytic residues are represented by black arrows. The three conserved motifs are highlighted by red boxes. The three conserved Cys are marked by triangles. (**E**) Conserved domain analysis of CsGPX proteins. TRX: Thioredoxin domain. Bar diagrams illustrate domain length and positions in each protein. (**F**) Predicted tertiary structures of CsGPX proteins generated using AlphaFold3. All proteins exhibit a well-resolved TRX fold in dark blue while N-terminal regions show intrinsically disordered or flexible conformations (low confidence in orange).

The close proximity of CsGPX2 and CsGPX4 on chromosome 5 suggests a possible duplication event. The CsGPX proteins range from 170 to 245 amino acids, corresponding to predicted molecular weights of approximately 19.3-27 kDa. CsGPX4 is the smallest GPX protein in *C. sinensis*. All CsGPXs have negative GRAVY values, indicating hydrophilic characteristics consistent with GPX families in other plant species (Khan et al., 2024). Interestingly, CsGPX4 displays an acidic predicted isoelectric point (pI= 4.76), in contract to the basic pI values (>79) observed for other CsGPX proteins. In addition, CsGPXs were predicted to be localized in the endoplasmic reticulum (ER), mitochondria, plastids, and cytoplasm (**Supplementary Table 1**).

On average, CsGPXs share approximately 50-75% sequence similarity (**Fig. 1C**). Multiple sequence alignment revealed a conserved characteristic of plant GPX proteins, including three conserved motifs VNAS[R/K/Q]CG, LAFPCNQF, and WNF(S/T)KF. The catalytic tetrad (cysteine (C), glutamine (Q), tryptophan (W), and asparagine (N)) of GPX was observed (Tosatto et al., 2008). The three conserved Cys residues in the active sites of plant GPXs were also present in citrus GPXs (**Fig. 1D**). All four CsGPX proteins contain a conserved TRX-like fold (**Fig. 1E**), consistent with previous reports that plant GPXs preferentially utilize TRX rather than glutathione as a reducing agent (Herbette et al., 2007b; Trenz and Margis-Pinheiro, 2025). AlphaFold3 structural predictions showed a well-resolved catalytic core with high confidence for the CsGPX proteins, whereas the N-terminal regions exhibited lower confidence scores (**Fig.1F**).

Among the four CsGPX proteins, CsGPX4 shares the highest similarity with AtGPX8 (**Fig 1A**; Fig **S1A**). Structural comparison further revealed a strong similarity between CsGPX4 and AtGPX8, with superimposition of the predicted models yielding a root mean square deviation (RMSD) of 0.364 Å (Fig. **S1B**). Given that overexpression of AtGPX8 was reported to enhance oxidative stress tolerance (Gaber et al., 2012), we investigated the post-translational regulation and functional dynamics of CsGPX4 in woody crops

### Overexpression of CsGPX4 in *C. sinensis* leads to oxidative stress phenotypes

We generated two stable *C. sinensis* lines overexpressing *CsGPX4* (GPX4-OX) driven by the cauliflower mosaic virus (CaMV) 35s promoter via *Agrobacterium*-mediated transformation (**Fig. 2A,B**). Transgenic shoots were identified by GFP fluorescence and downstream experiments. Surprisingly, both GPX4-OX lines exhibited oxidative stress phenotypes, including chlorosis and reduced growth, compared to the empty vector (EV) control plants (**Fig. 2B**). To determine whether CsGPX4 protein accumulated in the transgenic lines, immunoblot analysis was performed using an anti-HA antibody. Instead of a full-length CsGPX4-3xHA protein of 22.6 kDa, a prominent band approximately 11 kDa was detected in both GPX4-OX lines (**Fig. 2C**), suggesting that CsGPX4 underwent a post-translational truncation in citrus. PCR amplification with cDNA of the transgenic line as template produced the expected 505 bp full length CsGPX4 fragment in our CsGPX-OX lines (**Fig. 2D**). Sequencing of the amplified fragment eliminated the possibility of mutation of *CsGPX4* in the *GPX4*-OX lines indicating no variation in transcript, suggesting this to be post-translational modification (Fig. **S2**).

**Figure 2.**
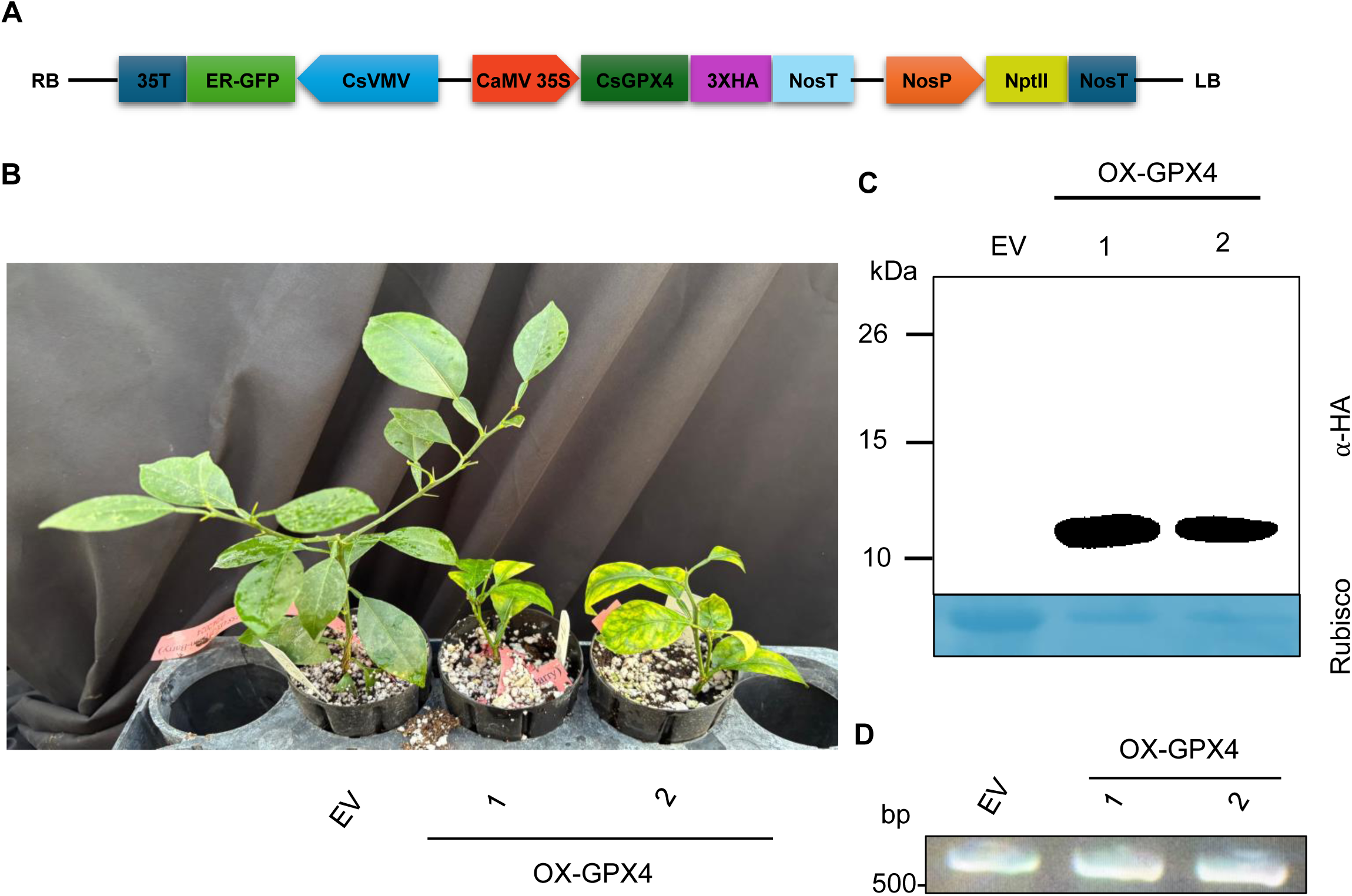
Verification of GPX4 overexpression and evidence of post-translational truncation in transgenic *C. sinensis* lines. **(A)** Schematic representation of plasmid constructs used for *CsGPX4* overexpression. (A) LB and RB, the left and right borders of the T-DNA region; CsVMV, the cassava vein mosaic virus promoter; GFP, green fluorescent protein; 35T, the cauliflower mosaic virus 35S terminator; NosP and NosT, the nopaline synthase gene promoter and its terminator; NptII, the coding sequence of neomycin phosphotransferase II. HA: Hemagglutinin tag. (**B**) Phenotypic comparison of empty vector (EV) and *CsGPX4* overexpression (OX-CsGPX4) *C. sinensis* plants. The picture was taken 1 year after planting in soil. (**C**) Immunoblot detection of HA-tagged CsGPX4 using an anti-HA antibody. Rubisco was used as a loading control. (**D**) Reverse transcription PCR confirmation of *CsGPX4* transgene expression in the OX-CsGPX4 lines. cDNA synthesized from total RNA was amplified using CsGPX4-specific primers. Amplicons of identical size were observed in both OX and EV plant, confirming correct transcript expression. PCR products were resolved by agarose gel electrophoresis.

To further investigate cellular effects associated with *CsGPX4* overexpression, we examined leaf midrib cells using transmission electron microscopy (TEM). Cells from EV plants displayed normal cellular architecture with compact cytoplasm, intact membranes, and well-organized vacuoles. In contrast, cells from GPX4-OX plants exhibited pronounced structural abnormalities. At the whole-cell level, GPX4-OX cells showed irregular cell boundaries and reduced cytoplasmic density compared with EV controls (**Fig. 3A**). Vacuolar morphology was also disrupted, with deformities in the tonoplast and intracellular membranes (**Fig. 3B**). Cytoplasmic organization was further compromised by extensive vesiculation and disordered intracellular contents (**Fig. 3C**). These abnormalities were accompanied by fragmentation of membrane systems, loss of membrane continuity, and accumulation of electron-dense materials, indicating severe cellular stress in GPX4-OX plants but not in EV controls (**Fig. 3D**). These results demonstrate that overexpression of *CsGPX4* leads to accumulation of a truncated CsGPX4 protein and unexpectedly induces oxidative stress phenotypes in citrus.

**Figure 3.**
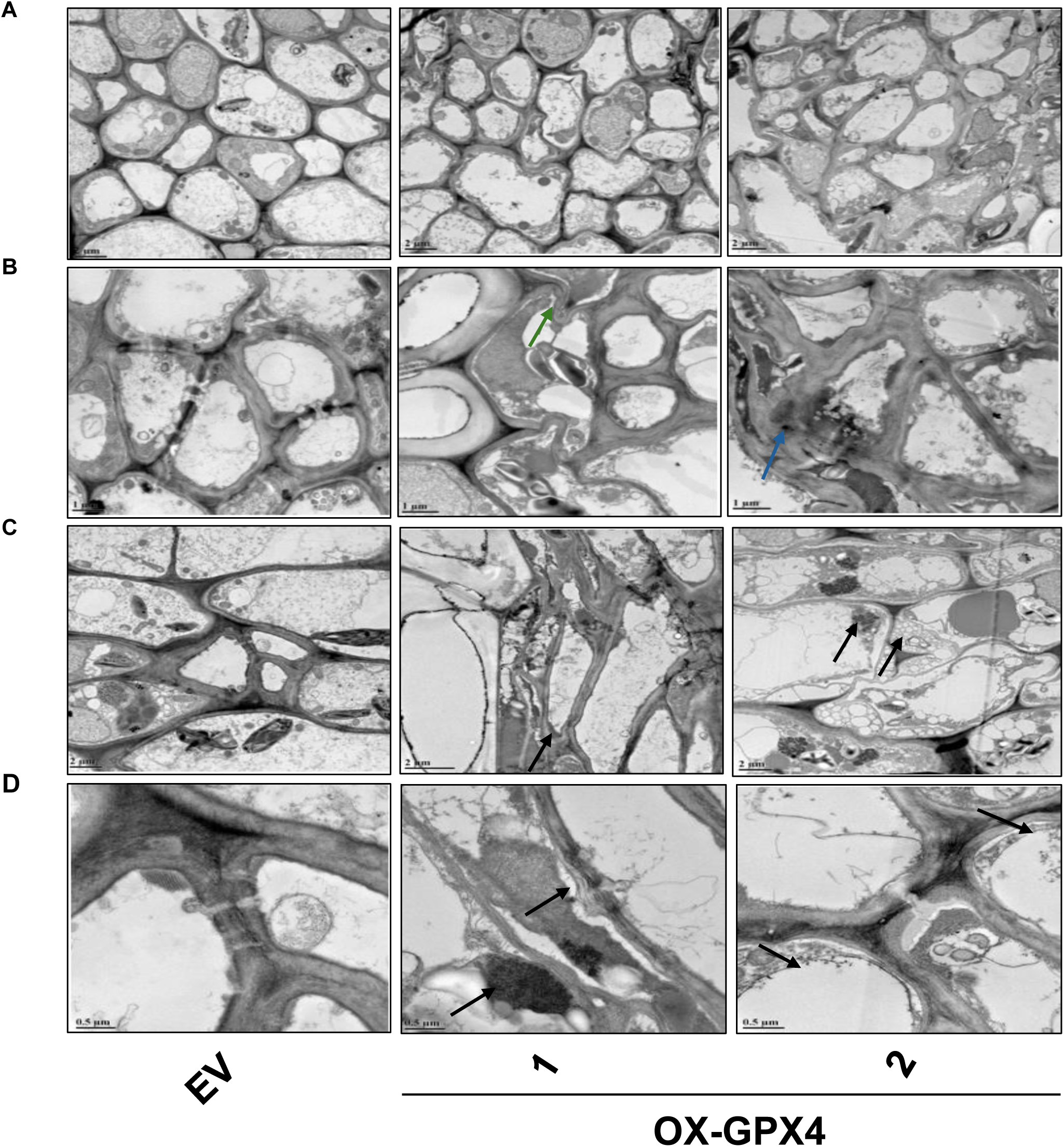
Transmission electron microscopy (TEM) analysis of midrib cells of *C. sinensis* leaves overexpressing *CsGPX4*. (**A**) Overall cell organization. CsGPX4-OX plant cells exhibit irregular boundaries and reduced cytoplasmic density compared to the empty vector (EV) transformed lines. (**B**) Vacuolar structure. CsGPX4-OX cells show pronounced tonoplast (green arrow) and membrane (blue arrow) deformation compared to EV. (**C**) Cytoplasmic organization. CsGPX4-OX cells display extensive vesiculation (arrow) and disorganization of intracellular contents (dashed arrow) compared to EV. (**D**) Membrane system integrity. OX lines show fragmented membranes, loss of structural continuity, and accumulation of electron-dense material (arrow), with more severe defects observed in OX-GPX4-2. Scale bars: (**A**) = 2 µm; (**B**) = 1 µm; (**C**) = 2 µm; (**D**) = 0.5 µm.

### CsGPX4 undergoes citrus-specific truncation and altered subcellular localization in *C. sinensis*

To determine whether the observed CsGPX4 truncation resulted from T-DNA insertion effects, whole-genome sequencing (WGS) was performed to identify the insertion site of a GPX4-OX line. The T-DNA was located at position chr2:1,646,389 within the UTR of the *SWEET2* gene (Fig. **S3**). Since mutation of *SWEET2* has no negative effects in growth or development of plants (Chen et al., 2015), it is unlikely that the abnormal phenotypes of the GPX4-OX line resulted from insertion in the UTR of *SWEET2*. To further investigate whether CsGPX4 truncation is host-dependent, we used the CsGPX4-3XHA plasmid (Fig. **S4A**) to transiently express *C. sinensis* and *N. benthamiana*. Immunoblot analysis showed that CsGPX4 accumulated as a truncated protein of approximately 11 kDa in *C. sinensis*, whereas a full-length protein of 22.6 kDa was detected in *N. benthamiana* (**Fig. 4A**). Transiently expressed *CsGPX4* in *C. sinensis and N. benthamiana* was purified and was run on the same SDS-PAGE gel (Fig. **S5**). These results indicate that CsGPX4 truncation occurs specifically in *C. sinensis*

**Figure 4.**
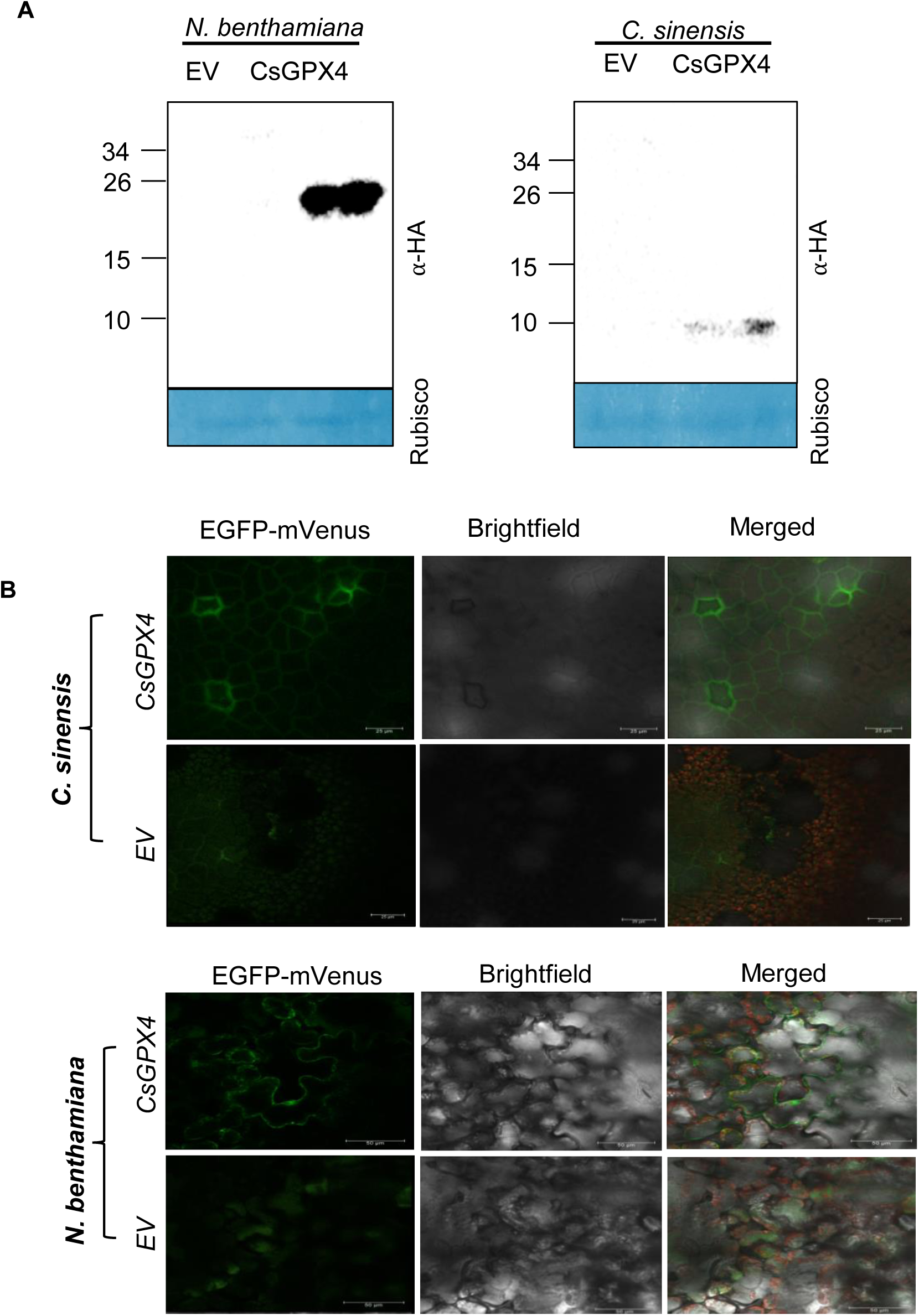
Host specific processing and subcellular localization of CsGPX4 in *C. sinensis* and *N. benthamiana*. (**A**) Immunoblot analysis of CsGPX4-3HA transiently expressed in *C. sinensis* and *N. benthamiana.* A truncated 11 kDa protein was detected in *C. sinensis* and a full-length protein of 22.5 kDa was detected in *N. benthamiana*. Rubisco was used as a loading control. (**B**) Subcellular localization of CsGPX4 in *C. sinensis* and *N. benthamiana.* Confocal microscopy of leaf epidermis expressing CsGPX4-EGFP-mVenus. Fluorescence displays a strong membrane-associated pattern in *C. sinensis*. Fluorescence displays GPX4 localization in cytoplasm and nucleus in *N. benthamiana*. Scale bars: *C. sinensis* = 25 µm; *N. benthamiana* = 50 µm.

Next, we examined whether truncation affects subcellular localization. We generated a construct where CsGPX4 was fused to mVenus (Fig. **S4B**). CsGPX4 was then transiently expressed in *C. sinensis* and *N. benthamiana* leaves. In *N. benthamiana*, CsGPX4 showed a cytoplasmic and nuclear localization pattern, consistent with previous studies in line with what was shown earlier (Gaber et al., 2012). In contrast, truncated CsGPX4 in *C. sinensis* exhibited membrane-associated localization (**Fig. 4B**, Fig**. S6**), suggesting that truncation alters the cellular distribution of the protein. These results demonstrate that CsGPX4 undergoes host-specific truncation in citrus, leading to altered subcellular localization.

### Truncation of CsGPX4 defines a cleavage site that impairs enzymatic activity

To determine the nature of this truncated proteoform, Matrix-Assisted Laser Desorption/Ionization-Time-Of-Flight Mass Spectrometry (MALDI-TOF) was performed on immunopurified CsGPX4 from citrus tissues. Masses were detected at approximately from 10,148.260 to 10,654.116 m/z, which validates that CsGPX4 was truncated when overexpressed in *C. sinensis* (**Fig. 5A**). Based on the observed masses with the subtraction of 3xHA tag length, the predicted cleavage region likely occurs within residues between L115-K117.

**Figure 5.**
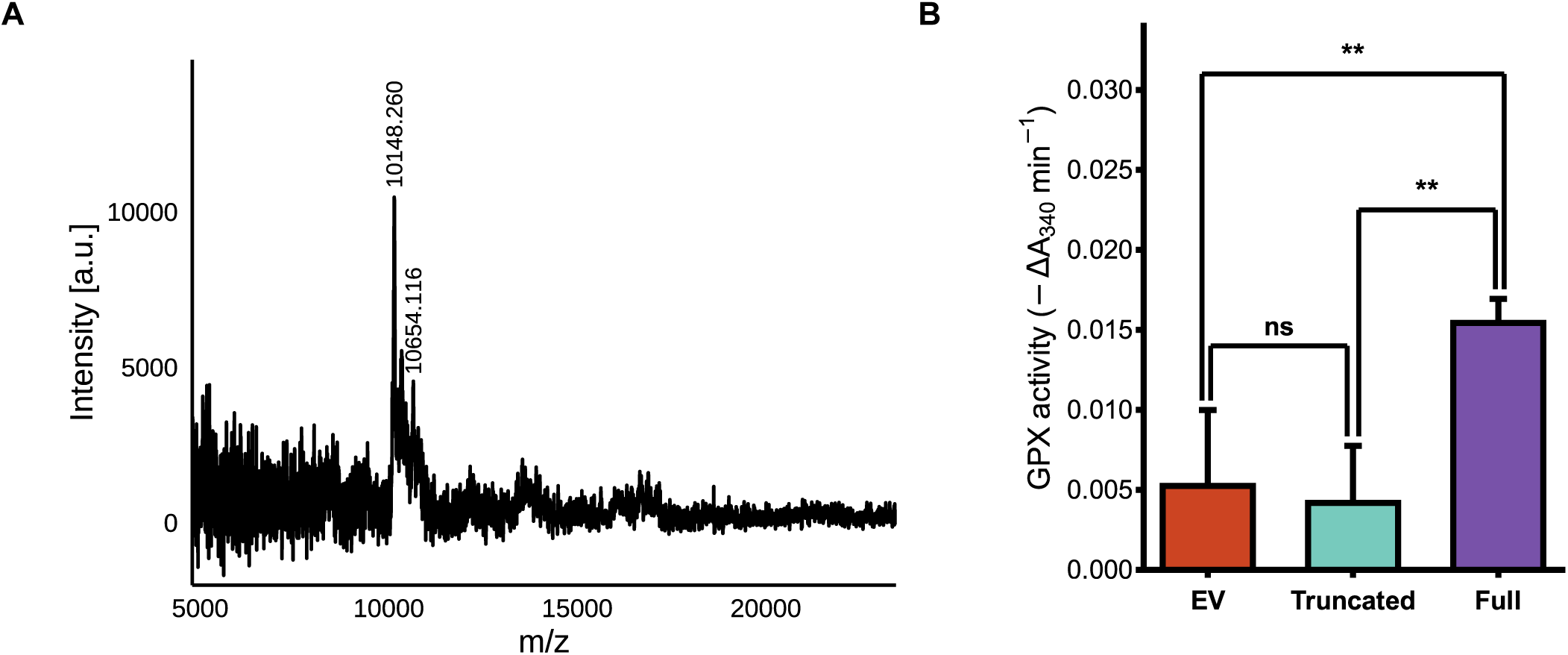
Predicted cleavage site impairs enzymatic activity of CsGPX4. (**A**) MALDI-TOF analysis reveals truncated CsGPX4 proteoform in CsGPX4-OX lines. MALDI-TOF mass spectrum of immunopurified CsGPX4 from citrus leaf tissues expressing HA-tagged CsGPX4 ranging (m/z) 6000-22000. Two prominent peaks were observed at m/z 10,548.260-10,654.116. Both masses are substantially lower than the predicted full-length protein (22.5 kDa). Intensity values are presented in arbitrary units (a.u.) scales at x104. (**B**) GPX enzyme activity assay of purified recombinant full CsGPX4 (Full), truncated CsGPX4 (Truncated), and empty vector expressed in E. coli. Initial linear reaction rates were calculated from the steady-state phase of the reaction of 1-5 minutes using absorbance of 340 nm per minute (ΔA320 min^−1^). Four biological replicates were used and statistical significance was determined using ANOVA followed by Tukey’s HSD. *P <0.05; **P <0.01; ns= not significant

To assess the functional consequences of this truncation, full-length and predicted cleavage site L115 proteins were designed using p15tv-L vector (Fig. **S5C**) (Pagliai et al., 2010) and expressed and purified in *Escherichia coli*. The p15v-L-CsGPX4 full and truncated plasmid was fused with a N-terminal 6xHis tag. Cell lysate and purified protein of CsGPX4-Full and CsGPX4-Truncated were ran on an SDS-page gel (**Fig. S7**). GPX activity was then measured using a coupled assay monitoring NADPH oxidation. The full-length CsGPX4 protein exhibited clear GPX activity, whereas the truncated CsGPX4 protein showed significantly reduced activity (**P<0.01) (**Fig. 5B**). Truncated CsGPX4 showed no significant difference between EV, indicating the truncation impairs enzymatic function. These results demonstrate that the proteoform of CsGPX4 showed reduced GPX enzyme activity compared to the full CsGPX4 protein.

### CsGPX4 overexpression disrupts intracellular ROS balance and reprograms the proteome in *C. sinensis*

To determine whether *CsGPX4* overexpression affects ROS accumulation in citrus, we performed ROS assays of intracellular and extracellular levels. Intracellular ROS levels were measured using the fluorescent probe H_2_DCFDA. GPX4-OX levels exhibited a significant increase in intracellular ROS levels compared to the empty vector control (EV) (*p* < 0.05), indicating that CsGPX4-OX disrupts intracellular ROS homeostasis (**Fig. 6A**). To evaluate H_2_O_2_ levels, extracellular ROS accumulation was quantified using Amplex Red and potassium iodide (KI) assays. KI is used as a reducing agent to detect peroxide (peracetic acid, persulfate, and H_2_O_2_) concentrations (Reichert et al., 1939; Sully and Williams, 1962; Klassen et al., 1994; Dai et al., 2021). Amplex red is a highly sensitive assay for H_2_O_2_ quantification that detects resorufin generated through the HRP-catalyzed reaction with H_2_O_2_ (Wang et al., 2017). Both assays showed no significant differences in extracellular H_2_O_2_ levels between *GPX4*-OX and EV plants (**Fig. 6B**, Fig. **S8**). These results showed CsGPX4 regulates intracellular ROS and disrupts its balance, rather than extracellular peroxide accumulation directly.

**Figure 6.**
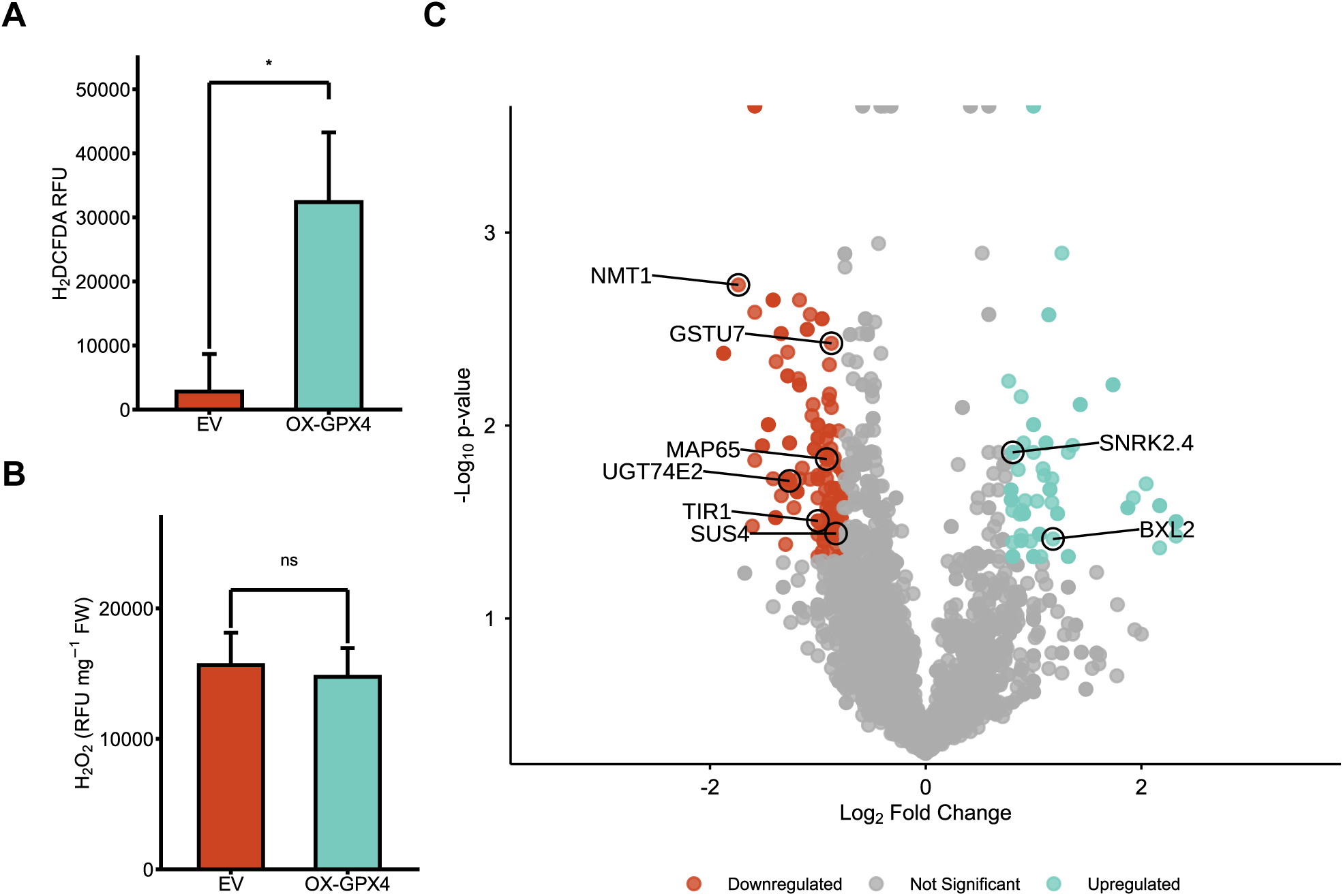
Quantification of ROS levels and proteomic analysis of the CsGPX4-OX lines and wild type. (**A**) Intracellular ROS levels measured using the fluorescent probe H_2_DCFDA. Three biological replicates were used and statistical significance was determined using students t-test (*P <0.05) (**B**) Quantification of extracellular H_2_O_2_ using the Amplex Red assays. Four biological replicates were used with EV having one removed due to technical error, student’s t test (ns) (**C**) Volcano plot showing differently expressed proteins in CsGPX4-OX citrus leaves relative to EV controls. Significantly downregulated proteins (red) labeled include NMT1, GSTU7, MAP65, TIR1, UGT74E2, and SUS4. Significantly upregulated proteins (teal) labeled include SnRK2.4 and BXL2. Non-significant proteins (gray). Three biological replicates were used.

To assess broader cellular responses associated with *CsGPX4* overexpression and to determine any metabolic shifts occurring, we performed global proteomic profiling of GPX4-OX and EV leaves (**Fig. 6C**). Differentially expressed proteins were associated with growth, cytoskeletal organization, hormone signaling and redox detoxification including N-myristoyltransferase 1 (NMT1; Cs_ont_1g019730.1), glutathione S-transferase U7 (GSTU7; Cs_ont_9g010810.1), microtubule-associated protein 65 (MAP65; Cs_ont_0419670.1), transport inhibitor response 1 (TIR1; Cs_ont_2g00600.1), UDP-glycosyltransferase 74E2 (UGT74E2; Cs_ont8g025420.1), and sucrose synthase 4 (SUS4; Cs_ont_9g001770.2). The upregulated proteins included SNF1-Related Protein Kinase 2.4 (SnRK2.4; Cs_ont_1g021000.1), and beta-D-xylosidase 2 (BXL2; Cs_ont_9g001770.2). Additionally, we identified 150 downregulated proteins and 91 upregulated proteins that were significantly expressed. There were 16 significantly differentially downregulated GO biological processes and two significantly differentially upregulated GO biological processes. Pathway enrichment analysis using PANTHER identified a set of significantly downregulated proteins involved in developmental pathways, including root and shoot system development, alongside with broader multicellular organism and anatomical structure developmental processes. Upregulated PANTHER pathway showed enrichment for protein modifications and stress-response pathways (Fig. **S9**).

## Discussion

In this study, we performed a systemic characterization of *C. sinensis* GPX family, showing only four members (*CsGPX1–CsGPX4)*. Previous studies showed that *Arabidopsis* contains eight GPX genes (Rodriguez Milla et al., 2003), and tobacco has 14 *GPX* genes (Peng et al., 2024). The GPX family in *A. thaliana* has undergone expansion (Zou et al., 2009) via whole-genome duplication (WGD) events, , whereas citrus species appear to maintain a more compact GPX repertoire. *A. thaliana* has undergone multiple rounds of WGD, including α, β, and γ events while citrus species have experienced only a shared ancient whole-genome triplication (γ -WGT) and lack recent WGD events (Bowers et al., 2003; Nakandala et al., 2025). GPXs play important roles in detoxification of ROS in different compartments. For instance, GPXs were reported to function in chloroplasts, cytosol, ER, mitochondria, nucleus and membrane (Ferro et al., 2003; Rodriguez Milla et al., 2003; Chang et al., 2009; Gaber et al., 2012; Attacha et al., 2017). Despite the small amount of GPXs in citrus, they were predicted or observed to function in most of the known GPX locations including ER, mitochondrion, plastid, cytosol and nucleus. Nevertheless, it is intriguing why citrus contains so few GPX genes compared to other plant species.

Previous studies showed that overexpression of *GPX8* increases plant tolerance against stresses such as oxidative stress (Gaber et al., 2012; Peng et al., 2024). In *Salvia miltiorrhiza*, overexpression of *GPX5* enhanced drought and oxidative stress tolerance without causing negative effects on plant growth and development (Zhang et al., 2018). Furthermore, overexpression of *TaGPX3.2A* and *TaGPX3.4A* in wheat under salt and drought stress enhanced ROS scavenging capacity (Jiang et al., 2023). Unexpectedly, overexpression of *CsGPX4*, a homologue of *AtGPX8*, leads to phenotypes commonly associated with oxidative stress, including growth inhibition, chlorosis, elevated ROS levels, and widespread cellular disruption (Savchenko and Tikhonov, 2021). It is unlikely that the abnormal phenotypes are due to overexpression of functional CsGPX4 protein or interfere with the detoxification enzyme activity of CsGPX4 or disruption of the gene function of insertion site. Our WGS analysis showed that T-DNA insertion occurs in the UTR of *SWEET2* gene. Prior studies indicate that *SWEET2* knockout mutants in *A. thaliana* do not exhibit significant phenotypic alterations under normal growth conditions (Chen et al., 2015). The truncation of CsGPX4 in citrus as evidenced by Western blotting and MALDI-TOF analyses (**Fig. 2C; 4B; 5A**) leads to loss of function as demonstrated in the *in vitro* the enzyme assay. Overexpression of truncated proteins in plants frequently results in dominant-negative effects, where the truncated protein disrupts the function of the full-length proteins, can compete with full-length proteins for binding partners, interfere with signaling or transcriptional regulation. For instance, overexpression of truncated ERECTA protein in *Arabidopsis* disrupts signaling of the whole ERECTA gene family and leads to reduced vegetative growth (Villagarcia et al., 2012). GPX proteins are known to interact with other proteins and have signaling functions (Miao et al., 2006; Del Río, 2015), raising the possibility that the truncated CsGPX4 proteoform could interfere with normal cellular signaling or redox regulation. Whether the truncated CsGPX4 exerts dominant-negative effects remain to be experimentally verified. GPX family members frequently exhibit partially overlapping antioxidant functions, but the distinct localization patterns along with signaling role could limit complete function redundancy among citrus GPXs. Therefore, the accumulation of truncated CsGPX4 proteoform could disrupt cellular processes in a manner that cannot be fully compensated by other GPX family members. The altered membrane-associated localization compared to cytosolic and nuclear of full length could contribute to the disruption of redox homeostasis. Whether nativeCsGPX4 undergoes similar proteolytic processes under normal conditions remains a mystery. Based on LC-MS result for global proteomic, we could not identify the endogenous CsGPX4, which possibly could be due to the low abundance or the proteome consists of a large deletion which could be hard for peptide mapping. Furthermore, we were able to identify other CsGPXs in both the EV and CsGPX4 datasets which suggest that other GPXs are still active. Further studies examining the endogenous CsGPX4 regulation during oxidative stress or pathogen infection would be important to determine whether a similar proteoform could generate in native physiological conditions.

Proteomic profiling revealed downregulation of key genes involved in detoxification and cellular structure, including NMT1, GSTU7, MAP65, TIR1, UGT74E2, and SUS4, and upregulation of SnRK2.4 and BXL2, linking CsGPX4 overexpression misregulation to impaired antioxidant defense, cytoskeletal destabilization, and altered hormone and cell wall signaling. Specifically, NMT1 is essential for post-translational lipid modification; its knockout causes severe growth arrest post-germination (Pierre et al., 2007). GSTU7 is involved in detoxifying ROS and enhancing stress resilience (Ugalde et al., 2021). MAP65 facilitates microtubule bundling and axial cell expansion (Lucas et al., 2011). TIR1 is a core auxin receptor; loss of function impairs hormone-mediated growth (Ruegger et al., 1998). UGT74E2 modulates oxidative stress response and water stress tolerance (Tognetti et al., 2010). Lastly, SUS4 is a key to sucrose metabolism and cell wall biosynthesis in citrus (Wang et al., 2018). On the other hand, upregulated proteins in CsGPX4-OX include several stress-response regulators. For instance, SnRK2.4 is known to mediate ROS signaling, antioxidant gene expression (e.g., catalase, ascorbate), and salt stress tolerance (Fujii and Zhu, 2009; Szymańska et al., 2019). These changes in CsGPX4-OX plants indicate a disruption in core metabolic pathways as observed in oxidative stress response (Savchenko and Tikhonov, 2021), likely contributing to the observed growth inhibition, microtubule instability, and reduced redox buffering in CsGPX4-OX plants.

In conclusion, our study identifies CsGPX4 as a critical regulator of redox balance in citrus based on the analysis of the overexpression of CsGPX4 and the consequences (**Fig. 6**). How overexpression of *CsGPX4* leads to its truncation in *C. sinensis* but not in *N. benthamiana* remains to be explored. The overexpression of CsGPX4 may perturb cellular redox homeostasis, potentially triggering compensatory post-translational regulation, including protein destabilization, or proteolytic cleavage. More broadly, our results position citrus as a valuable system for studying how post-translational mechanisms shape redox biology in perennial crops. The selective truncation of CsGPX4 underscores how long-lived plants may employ unique proteolytic or regulatory strategies to balance antioxidant activity with redox signaling. These findings have important implications for antioxidant engineering and underscore the need to consider host-specific regulation and protein stability when implementing GPX overexpression strategies.

## Materials and Methods

### Computational analysis of GPX orthologs

GPX homologs were identified from the *Arabidopsis* TAIR database and searched against the Citrus sinensis genome via BLAST using the Citrus Genome Database (https://www.citrusgenomedb.org/). Full-length GPX sequences from Arabidopsis, Citrus species were aligned using MAFFT and trimmed with Gblocks on NGPhylogeny.fr. Maximum likelihood trees were constructed in MEGA 12 (Kumar et al., 2024) with 1,000 bootstrap replicates. Phylogenetic tree was uploaded to iTOL (https://itol.embl.de/). Amino acid sequences of *C. sinensis* and Arabidopsis were uploaded to Jalview for comparison GPX proteins were domain searched using NCBI Conserved Domain Database (CDD). Identified GPX proteins were chromosome-mapped using MG2C (Chao et al., 2021). Multiple sequence alignments were performed with MUSCLE and visualized in Jalview (Waterhouse et al., 2009). Protein structure predictions for *C. sinensis* GPXs and AtGPX8 were performed using AlphaFold 3 (Jumper et al., 2021) with confidence visualized via pLDDT scores.

### Vector construction

The coding sequence of *CsGPX4* was amplified from *C. sinensis* cDNA using primers incorporating XbaI and KpnI restriction sites (**Supplementary Table. 2**).). The PCR product was gel-purified using the Monarch DNA Gel Extraction Kit and cloned into the binary vector RCsVMV-erGFP-pCAMBIA-1380N-35S-BXKES (Fig. **S2A**), previously digested with XbaI and KpnI (New England Biolabs), using the In-Fusion cloning system (Takara Bio). The resulting construct included a C-terminal 3HA tag and GFP under control of the CaMV 35S promoter.

The constructs pCAMBIA1380-35S-mVenus (Fig. **S2B**) for subcellular localization of *CsGPX4* and pCAMBIA1300S-35S-L279 (**S2A**) for transient expression (Wang et al., 2017) of *CsGPX4* and *AtGPX8* were similarly assembled, but SacI was used for digestion. Plasmid P15TV-L (Fig. S4B), digested with BseRI (Genbank accession EF456735) was used for protein purification for GPX enzyme activity (**Supplementary Table. 2**) (Pagliai et al., 2010). Constructs were verified by Sanger sequencing.

### Plant transformation

*Agrobacterium tumefaciens* strain EHA105 carrying binary constructs was cultured to OD600 = 0.4–0.5 in MS liquid medium (pH 5.4). *C. sinensis* seeds were surface-sterilized with 50% sodium hypochlorite, plated on Murashige and Skoog (MS) medium, and kept in darkness for 6–8 weeks. Epicotyl segments were excised, soaked in *Agrobacterium*-containing medium for 20 min, dried on sterile filter paper, and transferred to co-cultivation medium containing 6-Benzylaminopurine (BA), Indole-3-acetic acid (IAA), 2,4-Dichlorophenoxyacetic acid (2,4-D), and acetosyringone. After three days in darkness, explants were transferred to selection medium with kanamycin and timentin and in the dark for a month. Emerging GFP-positive shoots were micrografted onto Carrizo rootstock and transferred to MS medium supplemented with IAA and IBA. Upon establishment, plants were moved to growth chambers with plant growth hormones and placed in green house to soil when new leaves form.

### Agroinfiltration assay

*C. sinensis* and *N. benthamiana* leaves were infiltrated using the strain *A. tumefaciens* EHA105 strain. *A. tumefaciens* carrying the plasmid. A. tumefaciens was suspended in 10 mM Magnesium Chloride (MgCl_2_), 10 mM 2-N-morpholino ethanesulfonic acid potassium salt (MES-K), pH 5.6 with 100 μM of acetosyringone. OD600 of 0.6 was obtained and was mixed with *A. tumefaciens* carry vector pCAMBIA1300S-P19 in a 1:1 ratio. Roughly six-week-old *N. benthamiana* was inoculated and kept in Plant Growth Chambers from Geneva Scientific (#E-36HO Williams Bay, Wisconsin, United States). Growth chamber for *N. benthamiana* had a temperature of 23°C under high humidity in 16h light/8 h dark. Leaves were collected two days later. *C. sinensis* leaves were inoculated in a climate controlled cool greenhouse with dim lighting. Leaves were collected based on (Acanda et al., 2021) involving day point.

### Confocal localization analysis

Leaves were imaged for EGFP fluorescence using Confocal Laser Scanning Microscopy TCS-SP5 from Leica (Mannheim, Germany, Europe). Samples were excited with 488 nm and detected using photomultiplier tubes (PMT) tuned to 493-540 nm for EGFP.

### Western blot analysis

*C. sinensis* and *N. benthamiana* leaves were collected and placed in liquid nitrogen and TissueLyser II (Cat. #85300) from Qiagen (Hilden, Germany, Europe) that was used to homogenize the plant tissue. Total protein was extracted using CelLyticTM P Cell Lysis Reagent (#C2360) and 1x Protease Inhibitor Cocktail (#P9599) from Sigma (Kawaski, Japan). Lysis was centrifuged at 11,000xg for 15 minutes at 4°C. Supernatant was taken out and process was repeated two more times. Laemmli Sample Buffer (#1610747) from Bio-Rad (Hercules, CA, United States) with 10% of Beta-mercaptoethanol (βME) was mixed with lysis in 1:3 ratio and boiled at 99°C for 10 minutes. The denatured proteins of CsGPX4 and EV were separated using a 12% sodium dodecyl sulfate polyacrylamide gel electrophoresis gel (SDS-PAGE).

Immunoblots were washed in Tris Buffered Saline with Tween 20 (TBST) with 5% Bovine serum albumin (BSA). Antibody used in this study was Anti-HA High Affinity (#11867423001) from Roche (Basal, Switzerland, Europe). To visualize the immunoblots, SuperSignalTM West Pico PLUS Chemiluminescent Substrate (#34580) from Thermo Fisher was used. Loading control was confirmed by Ponceau S staining (#P7170) from Sigma-Aldrich (St. Louis, Missouri, United States) of the Rubisco large subunit (RbcL).

### Whole-genome sequencing and T-DNA insertion site identification

To identify the genomic insertion site of the CsGPX4 overexpression construct, whole-genome resequencing was performed using genomic DNA extracted from a transgenic plant line. Genomic DNA was extracted using the Promega Genomic DNA Kit (#A1120, Madison, WI, USA) and paired-end libraries were prepared and sequenced on the Illumina platform to generate 150 bp paired-end reads. Raw sequencing reads were assessed for quality using FastQC (https://www.bioinformatics.babraham.ac.uk/projects/fastqc/) and Trimmomatic (Bolger et al., 2014). High-quality reads were aligned to the *C. sinensis* genome (Zhong et al., 2024) concatenated with the full CsGPX4 T-DNA plasmid sequence using BWA-MEM.

To determine the T-DNA insertion site, read pairs were mapped to the T-DNA plasmid and to the *C. sinensis* genome. Integrative Genomics Viewer (IGV) (Thorvaldsdóttir et al., 2013) was used to visualize junction reads, soft-clipping events, and mate-pair connections. BEDTools was used to intersect insertion with the *C. sinensis* genome annotation in GFF3 format to assess gene disruption location.

### TEM analysis

Small leaf sections were collected and fixed in cold 3% Gluteraldehyde in 0.1M Sorensen’s Phosphate buffer (SPBS), pH 6.8 for 24 hours. Tissues were then rinsed three times for 30 minutes in 0.1M Sorensen’s Phosphate Buffer. Tissues were Osmicated in 1% osmium tetroxide for four hours room temperature. Tissues were then rinsed 3 times for one hour each in 0.1M SPBS. Tissues were dehydrated in acetone of concentrations 25%, 50%, 90% 100%, 100%, 100% for 20 minutes. Tissues were infiltrated in a 1:1 mixture of acetone wrapped in plastic overnight at RT. Infiltrated tissues were made in a 1:3 mixture of acetone and infiltrated again for six hours. Tissues were transferred to embedded molds, oriented and sat in RT until settled. The polymerize plastic was heated at 60°C for 12 hours.

The embedded tissue was placed on Epoxy resin blocks that were provided from University of Florida Interdisciplinary Center Electron Microscopy (ICBR-EM) Core for ultramicrotomy and TEM examination. Tissues were made into semi-thick resin sections and 500nm were collected onto super frost plus glass slides from Fisher Scientific (#22-037-246) and stained with toluidine blue. Ultrathin resin sections of 100nm were collected on carbon coated Formvar 2×1 mm nickle slot grids-TB from Electron Microscopy Sciences (EMS) (Hatfield, PA, United States). Ultrathin resin sections were treated with 1% meta periodate for 15 minutes and washed with H_2_O. Tissues were air dried and then post-stained with 2% aqueous uranyl acetate and citrate from EMS. Sections were then examined with a Field Electron and Ion (FEI) Teecnai G2 Spirit Twin TEM (Hillsboro, OR, United States) and Digital Micrograph Software Gatan (Pleasanton, CA, United States).

### H_2_O_2_ quantification

Extracellular H_2_O_2_ concentrations using KI method were measured based on (Kumar et al., 2011). Protocol was modification to fit onto a 96-well plate. Infiltrated *C. sinensis* tissues were weighed and ground and 1:4 of 0.1% (w/v) of trichloroacetic acid (TCA) and 5 µL of samples or standard was mixed with 28.25 µL of 1.0 phosphate buffer (PBS) at a pH of 7.0 and 16.25 µL of 1.0 potassium iodide (KI) solution. Plates were incubated in room temperature in the dark for five minutes. Absorbance of 290 nm was measured for the oxidized product. H_2_O_2_ concentrations were then calculated using the standard curve with H_2_O_2_ concentration and fresh weight measured in µmol/g. Three biological replicates were used. Additionally for extracellular H_2_O_2_ concentrations, Amplex^TM^ Red Hydrogen/Peroxide Assay from Thermo Fisher (#A22188) was used following manufacturer’s instructions with minor modifications (Garnier et al., 2006). Leaf tissues were collected and placed in liquid nitrogen and homogenized with fresh PBS, 7.2 at a ratio of 1:10 (w/v). Samples were incubated based on manufactors protocol and stop agent was added after incubation. Fluorescence was measured were normalized to tissue fresh weight and expressed using Relative fluorescence units (RFU) per mg fresh weight (RFU mg^−1^ FW). Four biologicals were used with EV having one removed due to technical error. Student t-test was used for statistical analysis.

For H_2_DCFDA Fluorescence Intensity, H_2_DCFDA stained was used from *C. sinensis* infiltrated with the following protocol with minor adjustments (Kapur et al., 2023) and manufactures instructions from Invitrogen (Eugene, OR, United States). Leaf size of 25 mm^2^ were collected and cleaned using distilled H_2_O and incubated in 10 µM H2DCFDA from Thermo Fisher (#D399) and were vacuum infiltrated for five minutes. Samples were washed using distilled H_2_O. Fresh PBS, pH 7.2 was added and incubated in the dark for 30 minutes. Fluorescence was measured using BMG LABTECH VANTAstar were read at a fluorescence of Ex/Em= 485/535 nm in end point mode and analysis was performed and RFU was calculated based subtraction of background autofluorescence. (Aimonen et al., 2022). Three biological replicates were used. Student t-test was used for statistical analysis.

### Shotgun proteomics

Infiltrated *C. sinensis* leaves were collected and placed in liquid nitrogen and using the TissueLyser II to homogenize. Total protein was extracted using 50 mM Tris-HCL, pH 8.0, 0.01% n-dodecyl- β-maltoside (DDM), and 1x of Plant Proteaserrest^TM^ Protease Inhibitor Cocktail (#786) from G-BioSciences (St. Louis, MO, United States). Lysis preparation had same protocol was western blot method. An 12% SDS-PAGE gel was made with 50 µg of lysis protein loaded with three biological replicates of CsGPX4 and EV and was ran very briefly. Each lane was excised for in-gel trypsin digestion using Trypsin. Trypsin (#V5280) from Promega (Madison, WI, United States) was used for digestion; maufactor protocol was applied. The peptides were desalted using Pierce^TM^ C18 Spin Tips & Columns (#87784) from Thermo Fisher following manufacturer’s manual and samples were analyzed using a Thermo Orbitrap Exploris 240 combined Vanquish Neo UHPLC with a 120-minute LC gradient (Jiang et al., 2025). Mass spectrometry data were in data-dependent acquisition (DDA) mode. Full MS scans (MS1) were collected over an m/z range of 375-1475 at a resolution of 120,000. MS/MS fragmentation was performed at a resolution of 15,000, targeting the most abundant precursor ions within a 3-second duty cycle. The normalized AGC target is set to 300% for MS1, and a standard MS2 AGC target was applied using a normalized HCD collision energy of 30%. Dynamic exclusion was enabled with a 45-second exclusion duration. PatternLab V was used for data analysis with extracted ion chromatograms from protein expression profiles (Santos et a., 2022). Protein expression changes were evaluated using extracted-ion chromatograms (XIC). Briefly, two-sample Student’s t-test was used for differential analysis with p cutoff being 0.05. To control false discovery rate, Banjamini-Hochberg FDR estimator was included in the TFold module. F-stringency, L-stringency and q-value were set to 0.1, 0, and 0.05 as default settings. Minimal absolute fold change for proteins with significance was consequently 1. Three biological replicates were used.

### MALDI-TOF MS

Total protein was processed as stated in Western blot methods section but used a motor and pestle. Pierce^TM^ Anti-HA Agarose (#26181) from Thermo Fisher was used for immunoprecipitation. Lysates were incubated rotating with Anti-HA Agarose slurry at 4°C for four hours. Reaction was washed using 50 mM Tris-HCL, pH 8.0. Proteins were eluted using elution buffer (0.1M Glycine, pH 2.0 and 1:10 1M Monoammonium Phosphate (ABC)). C18 Spin Tips & Columns (#87784) from Thermo Fisher was used for desalting following manufacturer’s instructions. Protein at 50 µL was concentrated at 10 uM.

For Matrix-assisted laser desorption/ionization time-of flight (MALDI-TOF), 1 µL was aliquot of purified protein was mixed 1:1 with saturated sinapinic acid (SA) matrix prepared in 70% CAN and 0.1% TFA. A 1 µL aliquot of this mixture was spotted onto a stainless steel MALDI target plate and air dried. Spectra were acquired on a Bruker autoflex maX MALDI-TOF (Billerica, Massachusetts, United States) mass spectrometer operating in linear positive ion mode over an m/z range of 5,000-22,000. Spectra were acquired using FlexControl and analyzed with PolyTools 2.0. Signal intensities were reported in arbitrary units (a.u.) scales ×10^4^.

### GPX enzyme activity assay

The CsGPX4-Full, CsGPX4-truncated, and EV constructs were expressed in *E.coli* BL21-Star (DE3) cells (Agilent Technologies, Santa Clara, CA). The cells were grown in LB medium at 37°C to reach an OD_600_=0.6. Expression was induced using 0.1 mM isopropyl β-D-1-thigalactopyranoside (IPTG) and incubated for 12 hours using 20°C. Cells were harvested and resuspended in binding buffer (500 mM NaCl, 5% glycerol, 50mM Tris pH 8;0, 5 mM imidazole, 0.5 mM TCEP) based on Pagliai et al., 2010. Cells were lysed and passed through French Press. The lysate cells were centrifuged at 17,000 rpm for 30 minutes at 4°C. Supernatant was stored in −80°C to run on SDS-Page gel for visualization and suspended in metal chelate affinity-column charged with Ni^2+^ (#PI88221) for one hour in 4°C rotating. Protein was washed and eluted using elution buffer (binding buffer with 250 mM Imidazole) (Pagliai et al., 2010). The purified protein was dialyzed using Snakeskin tubing (#88242) in dialysis buffer (10 mM Tris pH 8.0, 500 mM Nacl, 0.5 mM TCEP, and 2.5% glycerol) overnight. Protein was centrifuged at 12,000 x g for five minutes at 4°C. Concentration was determined using DeNovix DS-11 FX+ (Wilmington, Delaware, USA). Supernatant was used for GPX enzyme activity assay and used for SDS-PAGE Gel for visualization using 0.25%Coomassie Brilliant Blue R-250 powder (#1610400) dissolved in 40% H_2_O, 45% methanol and 7% glacial acetic acid.

GPX enzymatic activity was measured using spectrophotometric method (Drotar et al., 1985), but using the microplate reader modification (Halusková et al., 2009) (Agilent BioTek Synergy H1 Multimode Reader, Santa Clara, CA, USA). The reaction mixture (250 μL) containing 100 mM sodium phosphate buffer, pH 7.0, 1 mM EDTA, 2 mM glutathione, 0.1 U glutathione reductase, 0.2 mM nicotinamide dinucleotide phosphate (NADPH), and 4.8μg of purified protein. Reaction incubated for one minute and reactions were initiated with 1 mM of H_2_O_2_ The NADPH oxidation rate was measured at 340 nm over a period of five minutes, and calculation was based on the rate of absorbance (ΔA340 min⁻¹) decrease. Four biological replicates were used. Student t-test was used for statistical analysis.

## Author Contributions

S.B. and N.W. conceived the work. S.B., N.S.S, and N.W. and drafted the manuscript. S.B. also designed and preformed experiments. X.W. ran the CsGPX4 and EV samples in LC-MS for global proteomics.

## Acknowledgements

We thank Dr. Karen Kelley at University of Florida Interdisciplinary Center for Biotechnology Research for TEM expertise and Dr. Kari Basso for running our sample into the MALDI-TOF. Additionally, we would like to thank Dr. Zhonglin Mou for providing us the pCAMBIA1300. We would like to thank our funding sources of the Emergency Citrus Disease Research and Extension Program grants 2022-70029-38471 and 2021-67013-34588 and the U.S. Department of Agriculture’s National Institute of Food and Agriculture grant 2023-70029-41280 to Dr. Wang and 2024-33522-43698 to Dr. Nadakuduti. MALDI-TOF funding was provided from NIH grant award numbers S10 OD021758-01A1 and S10 OD030250-01A1

## Completing interests

All authors declare no conflict and competing financial interests.

## Data Availability

Whole genome resequencing data for CsGPX4 were deposited to NCBI under project ID (PRJNA1419271). Global proteomic data of CsGPX4 and EV are deposited in MassIVE repository (MSV000100757) and ProteomeXchange Consortium (PXD074185).

---

**Supplementary Figure 1. Comparison of the CsGPX4 and AtGPX8. (A) Multiple sequence alignment of CsGPX4 and AtGPX8 proteins generated with Clustal omega.** (B) AlphaFold 3-predicted tertiary structures of CsGPX4 (cyan) and AtGPX8 (tan). Superimposition (bottom) revealed a high conserved TRX-like fold, with a RMSD of 0.364 Å.

**Supplementary Figure 2. Sanger sequencing results of amplified cDNA fragment and alignment of CsGPX4-OX lines (GPX4-1;GPX4-2) and empty vector control (EV).**

**Supplementary Figure. 3 The T-DNA insertion site was identified through whole-genome sequencing and mapped to the 5′ untranslated region (5′ UTR) of *SWEET2* (Cs_ont_2g002980.3) on chromosome 2 (chr2:1,646,389 on the negative strand).** Arrow indication of T-DNA insertion site; Red: promoter region; blue: UTR; yellow: CDS.

**Supplementary Figure 4. Schematic representation of plasmid constructs used for *CsGPX4* overexpression in transient expression & protein purification.** (A) Schematic representation of the 3xHA-tagged CsGPX4 construct used for transient expression. RB and LB, right and left borders of the T-DNA region; CaMV 35s (enhanced), Cauliflower mosaic virus 35s enhanced promoter; 3xHA, triple hemagglutinin epitope tag; CaMV poly(A), cauliflower mosaic virus polyadenylation signal; Lac operator, Lac promoter, Catabolite Activator Protein binding site, and Lac fragment, bacterial lac regulatory elements. (B) Schematic representation of the plant expression vector used for subcellular localization assay. RB and LB, right to left borders of the T-DNA region; CaMV 35s, Cauliflower mosaic virus 35s promoter; mVenus, yellow fluorescent protein reporter; NosT, nopaline synthase terminator; 6xHis, 6 histidine affinity tag. (C) Schematic representation of the bacterial recombination expression vector used for CsGPX4 protein expression. T7 promoter and T7 terminator, bacteriophage T7 transcriptional regulatory elements; RBS, ribosome binding site; 6xHis, 6 histidine affinity purification tag.

**Supplementary Figure 5. Full immunoblot of CsGPX4 agroinfiltrated in *C. sinensis* (approximately: 11 kDa) and *N. benthamiana (*22.5 kDa).** Left: CsGPX4 in *N. benthamiana* in comparison of EV and negative control; Right: CsGPX4 in *C. sinensis* in comparison of EV and negative control. M: marker; Rubisco is serving as loading control.

**Supplementary Figure 6. Subcellular localization of CsGPX4 in *N. benthamiana in cytosol*.** Confocal laser scanning microscopy showing the localization of CsGPX4 fused to the mVenus reporter at the C-terminus.

**Supplementary Figure 7. SDS page of CsGPX4-Full and CsGPX4-Truncated of cell lysate (supernatant) and elution (purified protein). M: protein ladder.**

**Supplementary Figure 8. Quantification of extracellular H_2_O_2_ using the KI assays.** Three biological replicates were used.

**Supplementary Figure 9. PANTHER pathway enrichment analysis of upregulated (top) and downregulated (bottom) of significant proteins from global proteomics are shown.**

