## Supplementary Figures for "Citrus-specific truncation of CsGPX4 disrupts intracellular ROS homeostasis"

**<u>Supplementary Information</u>**

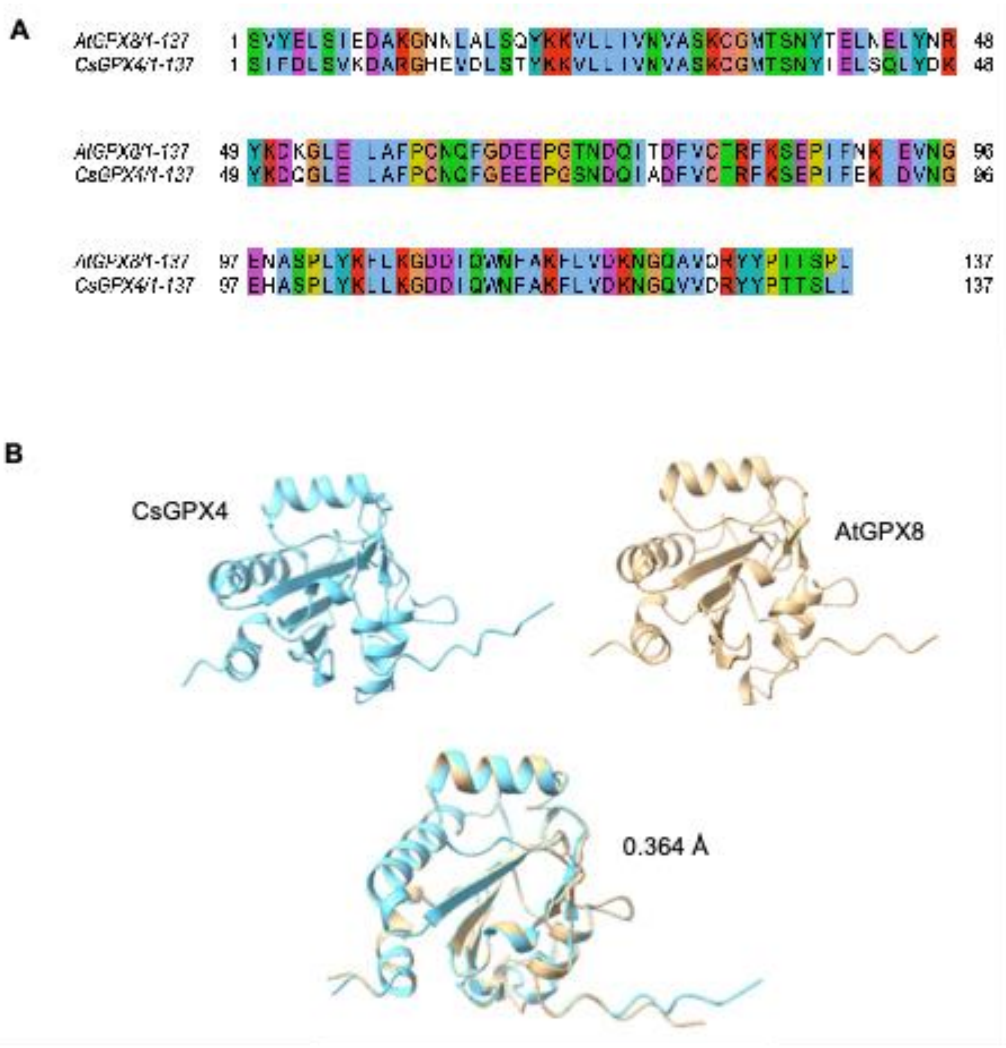


**Supplementary Figure 1. Comparison of the CsGPX4 and AtGPX8. (A)** Multiple sequence alignment of CsGPX4 and AtGPX8 proteins generated with Clustal omega. (**B**) AlphaFold 3-predicted tertiary structures of CsGPX4 (cyan) and AtGPX8 (tan). Superimposition (bottom) revealed a high conserved TRX-like fold, with a RMSD of 0.364 Å.

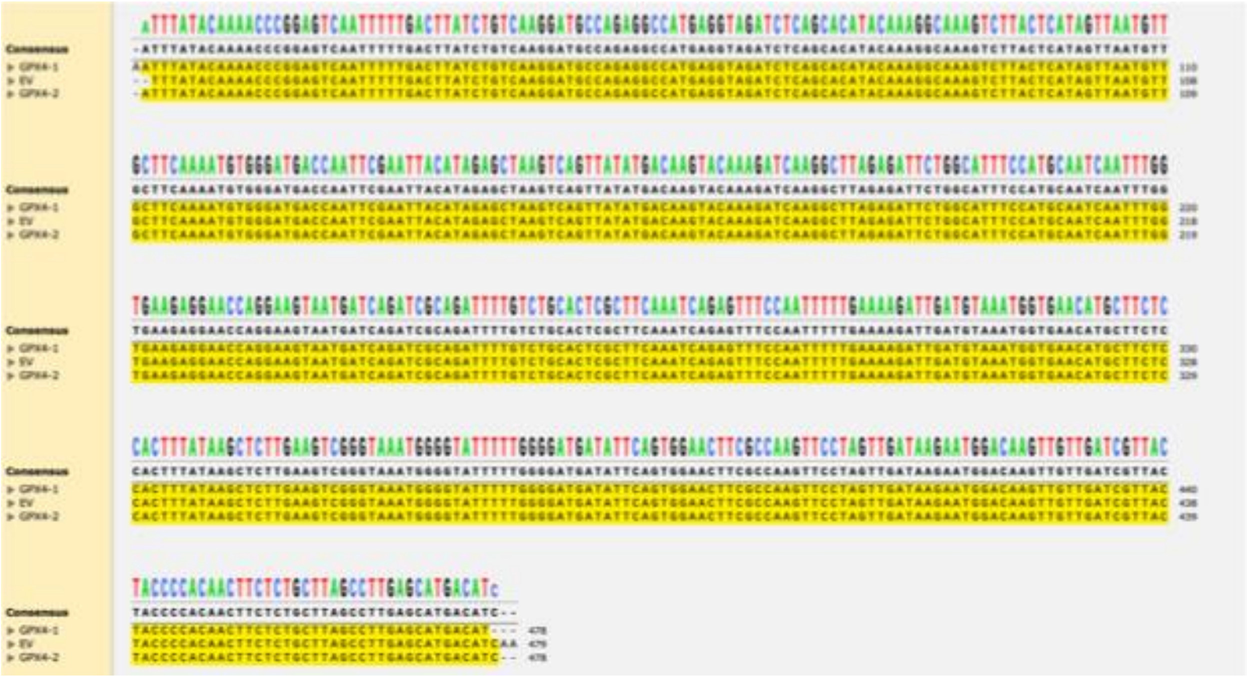


**Supplementary Figure 2. Sanger sequencing results of amplified cDNA fragment and alignment of GPX4-OX lines (GPX4-1;GPX4-2) and empty vector control (EV).**

**>Cs_ont_2g002980.3-gene_updownstream chr2:1,643,759-1,646,458(-) +/- 100 up/downstream bp**

ttcttctgagttccacttggagtctatcaaatatcccctccacgaaagttcctcggttccccagtattattaaattactatatattaaatatacg
acacacaaagacacagttgatctccagtctccaccacatatattattttagaaagggataatatttctaatatgcgttgatttgatgtgcctc
tgtttttttttattttatatatttatctaaaaagacaaaatcatttgaaaggcaaattcaaggccactagacataagaagctttaacagacac
ataacttcacaatttgacaaatgaatggctttaaatgttccttcaaggcatactacatatatttattcgtatatggtctttcacaagcaaacttt
aagttctgcacaaagctacaaattagcaaggcagaagaggggcagagagagtcggcattggtgttgaattgagtgatcagtgaatt
gcaagtgaagttttagggagatgtcttcagttgggatttcttcaatttactcaggttgtagtgttgcagctggcgttactggtaaatttttcatca
tcctacaatctcatctgctgcattattcttcttttaatatgcaagattgaaatcttatatgcttttttttttcctttttttggggcttaaagattgtacatc
taacttgttttctttattttggtcttgtttaaactgtttcaattgaagaagagtgacttggcttttgttgtttttttgttgccaagaacatgcagtgagga
aagaaaagttgaaaattcttgcttcgtagaatctttagcatgcaaagttccggtatctaaattaattgctttctagagtttctcacaaactata
tgtttcactttctctctatggtaaagaattgctctaatatagagaagtttacatgaattcccatttggtaaaatcggaactctagtacactgca
gtttttccaacactgtgatgatggtacatgtgattgcgatgaattgatcaaccttaattccagattataatttgttaattgcttcagtttctggaa
gctaaagtgaaagtattctaatactattttattgattgcagggaacatatttgcttttgtattgtttgtatcgcccatgtaagttgttgtttaataatc
tttccatttaaacaagaattatattgggaaaatgaaactccttgagcttatttcaatatttttctccctctttcagacctacattcaggagaatttt
gagaaacaagtcaacagaacaattctcaggattgccttacatctgttctctcttgaattgcttgattaccctttggtatggcATGCCTC
TTGTATCCCCTGGTATTATATTAGTCGCTACAGTGAATTCAGTTGGAGCAGTTTTCCAGTTG
ATTTATGTTAGCATCTTCATTTCATATGCTGAAAAAGCTATAAAGgtaattcttttatgcttgattggtcttgtt
attggaatgatgagaaattctgtcatatctcattaccatctttgaatgccttttccagCTGAAGATATCTGGATTATTGATA
GCAGTTTTTTTAGTATTTCTTGCCATAGTCTTCACGAGCATGGAAGTTTTTGACTCCAATGG
GCGGCGGCTTTTTGTTGGATATTTGAGTGTTGCTTCTCTCATTTCTATGTTTGCTTCGCCAT
TATTCATTATTgtaagctcatctggaactcaagcattccgacttttaagactacacatttccttacattcatatggttgcatgtatatttt
tatgcagAAGTTAGTCATTAAGACGAGGAGCGTTGAATTCATGCCCTTTTATCTTTCTCTTTCA
AATTTCTTGATGAGTCTCTCCTTCCTAGCTTATGGATTGTTCAAGGATGACCCTTTCATTTA
Tgtaagtactatctttcctgcagaaccattatagttaattctagattttcatgtaacattgacaagcagagtaggcatgacagacttaaaa
aaaaaaaaaaaaggaattgtacttttacagttaatgaatcagtggtgggctgaattgttgctgtttaactccaagtactatctttttgcttcgtt
tgtgtactcagaaatcaattcaaacaataactatgagtattgcatcagagaattttgattcatttcattatcattttcggatccagGTCCC
AAATGGAATTGGAACACTTCTGGGGATTGCTCAAGTGATGTTGTATTCCTACTATAGCACT
AAATCTGGTGAAGTCTCAAGACAACCCCTGATCGATTCATTTGCATAAattaatgtgaaaatcatgtgt
agcagagaagttagcgatcagcatctgtataatcagtaattcagtgacttgattattctaaagctcgaattcctttggaatctaggcgcttc
actacttgaggtctgaggcagcaaatgtaagactggtaaaatgctattttagactttccaaatcactcattaagcaacaagtaatcttata
tggagctacgtgcccttgttgaacgcagagacaagtctcaatcacacaacattgtccaggcttgctagctgatagatgggatgtcaagt
ggattttatttgagcttgatcccttctattgccaaatgccttgtcttattctttactgcgtccaaatcatttgaattaagtttatatggaagaagaat
cagatatctgatatatcgatgaatctatatcttatgctgctcttgtattagagcttcctcaagtgagatgtatattgagcaattatcagcataa
atgaaatttcaacactcagctattctgggtaactttttttgcatgataaatcttctgtccctgcagtaaatgtttatatcaaacaaaagttttagta
tgggaagaggtaattggtacaaagtgtagtttgttttaatgca

**Supplementary Figure 3. The T-DNA insertion site was identified through whole-genome sequencing and mapped to the 5′ untranslated region (5′ UTR) of *SWEET2* (Cs_ont_2g002980.3) on chromosome 2 (chr2:1,646,389 on the negative strand).** Arrow indication of T-DNA insertion site; Red: promoter region; blue: UTR; yellow: CDS.

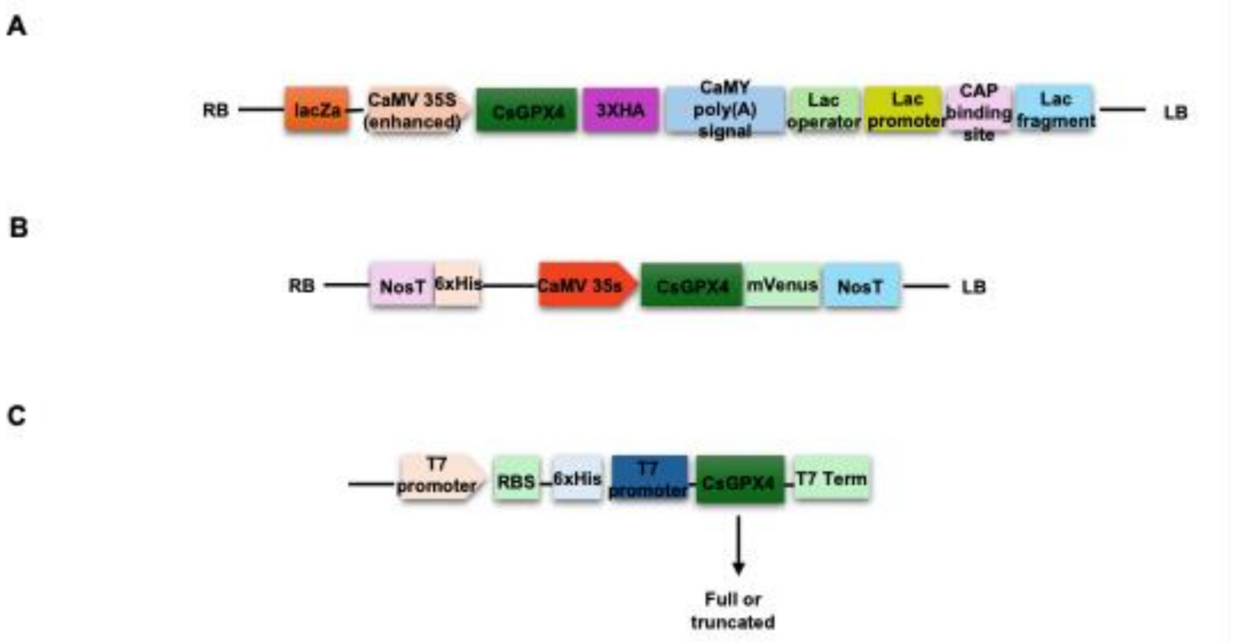


**Supplementary Figure 4. Schematic representation of plasmid constructs used for *CsGPX4* overexpression in transient expression & protein purification.** (A) Schematic representation of the 3xHA-tagged CsGPX4 construct used for transient expression. RB and LB, right and left borders of the T-DNA region; CaMV 35s (enhanced), Cauliflower mosaic virus 35s enhanced promoter; 3xHA, triple hemagglutinin epitope tag; CaMV poly(A), cauliflower mosaic virus polyadenylation signal; Lac operator, Lac promoter, Catabolite Activator Protein binding site, and Lac fragment, bacterial lac regulatory elements. (**B**) Schematic representation of the plant expression vector used for subcellular localization assay. RB and LB, right to left borders of the T-DNA region; CaMV 35s, Cauliflower mosaic virus 35s promoter; mVenus, yellow fluorescent protein reporter; NosT, nopaline synthase terminator; 6xHis, 6 histidine affinity tag. **(C)** Schematic representation of the bacterial recombination expression vector used for CsGPX4 protein expression. T7 promoter and T7 terminator, bacteriophage T7 transcriptional regulatory elements; RBS, ribosome binding site; 6xHis, 6 histidine affinity purification tag.

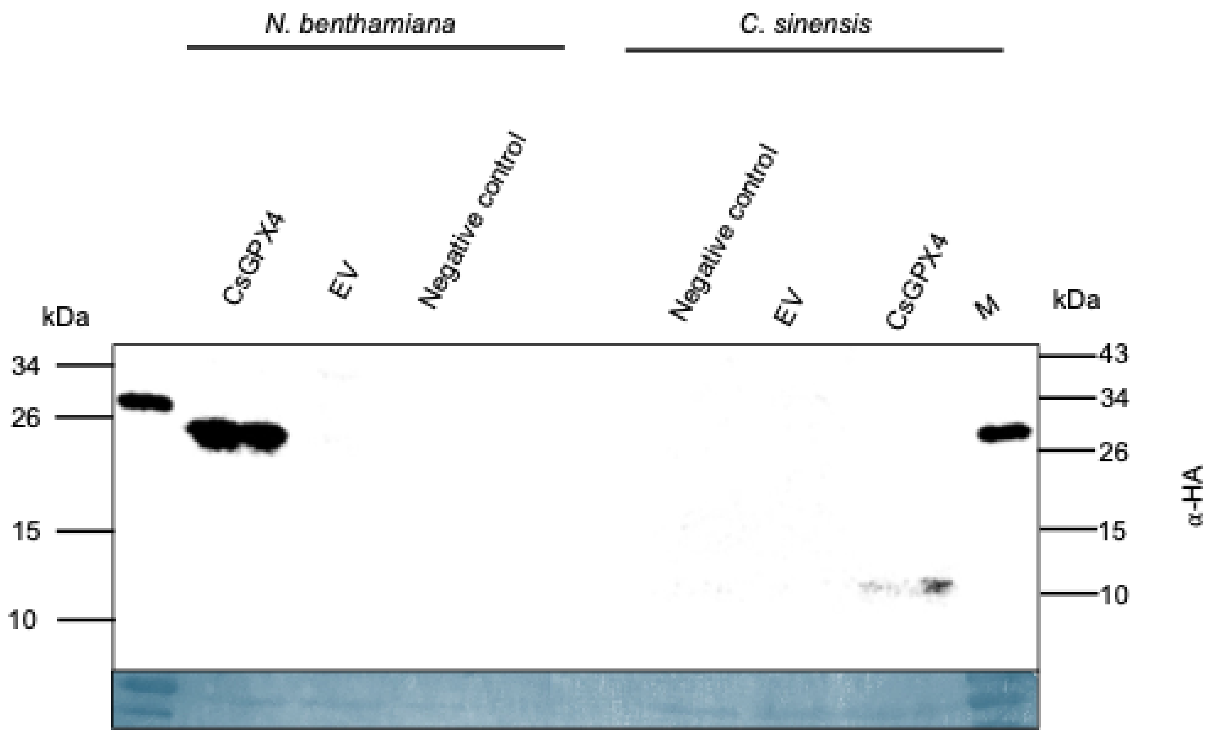


**Supplementary Figure 5. Full immunoblot of CsGPX4 agroinfiltrated in *C. sinensis* (approximately: 11 kDa) and *N. benthamiana (*22.6 kDa*).*** Left: CsGPX4 in *N. benthamiana* in comparison of EV and negative control; Right: CsGPX4 in *C. sinensis* in comparison of EV and negative control. M: marker; Rubisco is serving as loading control.

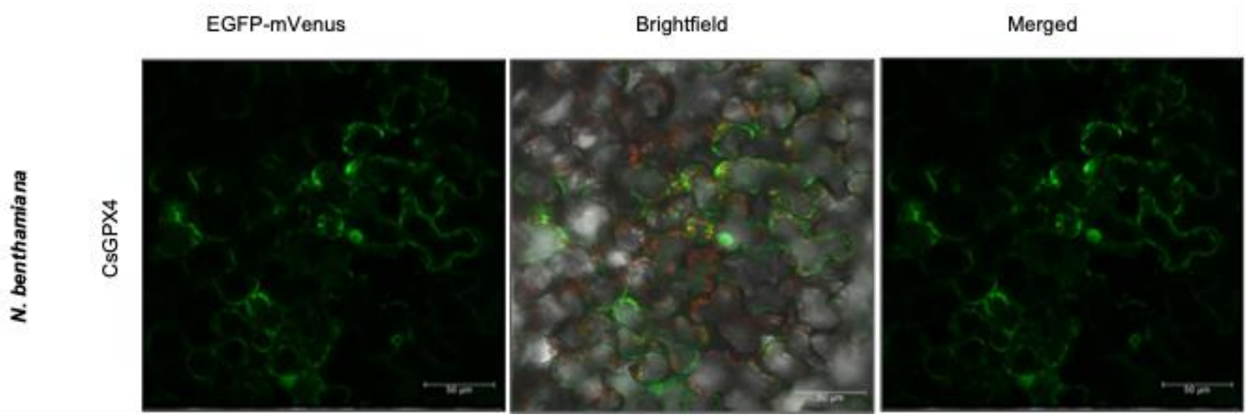


**Supplementary Figure 6. Subcellular localization of CsGPX4 in *N. benthamiana* cytosol.** Confocal laser scanning microscopy showing the localization of CsGPX4 fused to the mVenus reporter at the C-terminus.

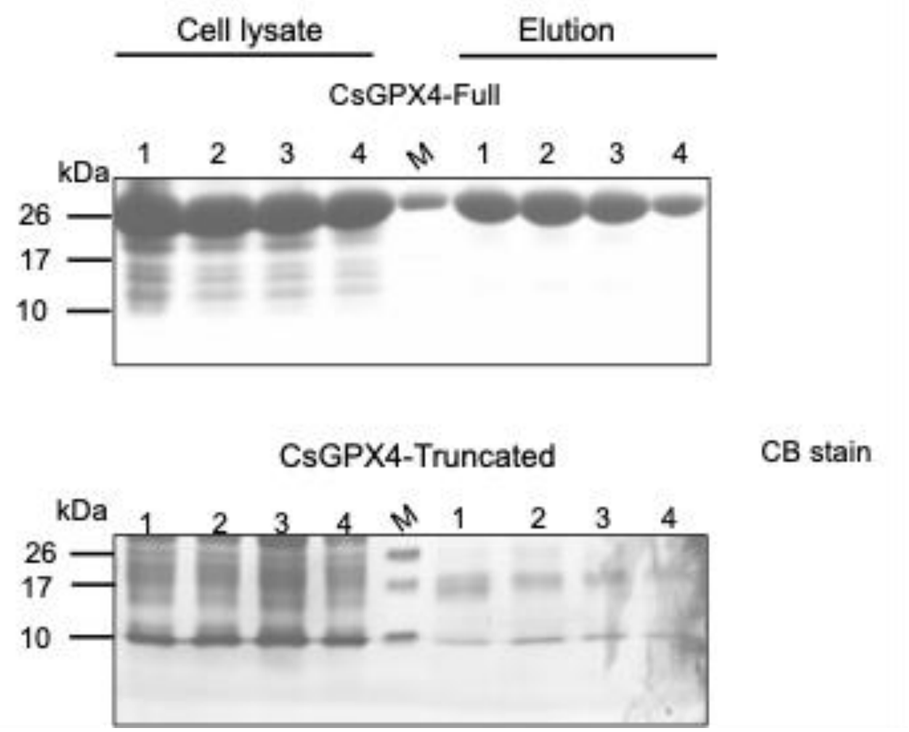


**Supplementary Figure 7. SDS-PAGE analysis of CsGPX4-Full and CsGPX4-Truncated of cell lysate (supernatant) and elution (purified protein).** M: protein ladder.

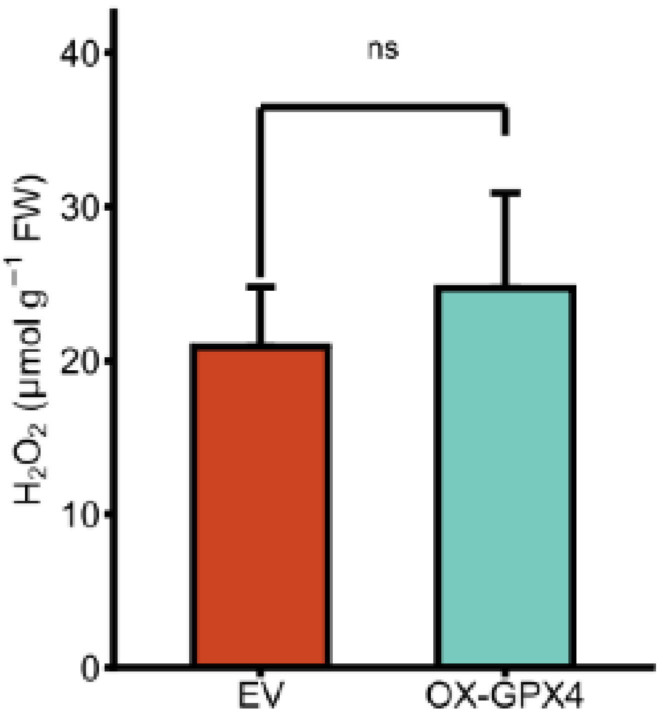


**Supplementary Figure 8. Quantification of extracellular $H_2O_2$ using the KI assays.** Three biological replicates were used. Student's t-test. ns: not significant $p < 0.05$.

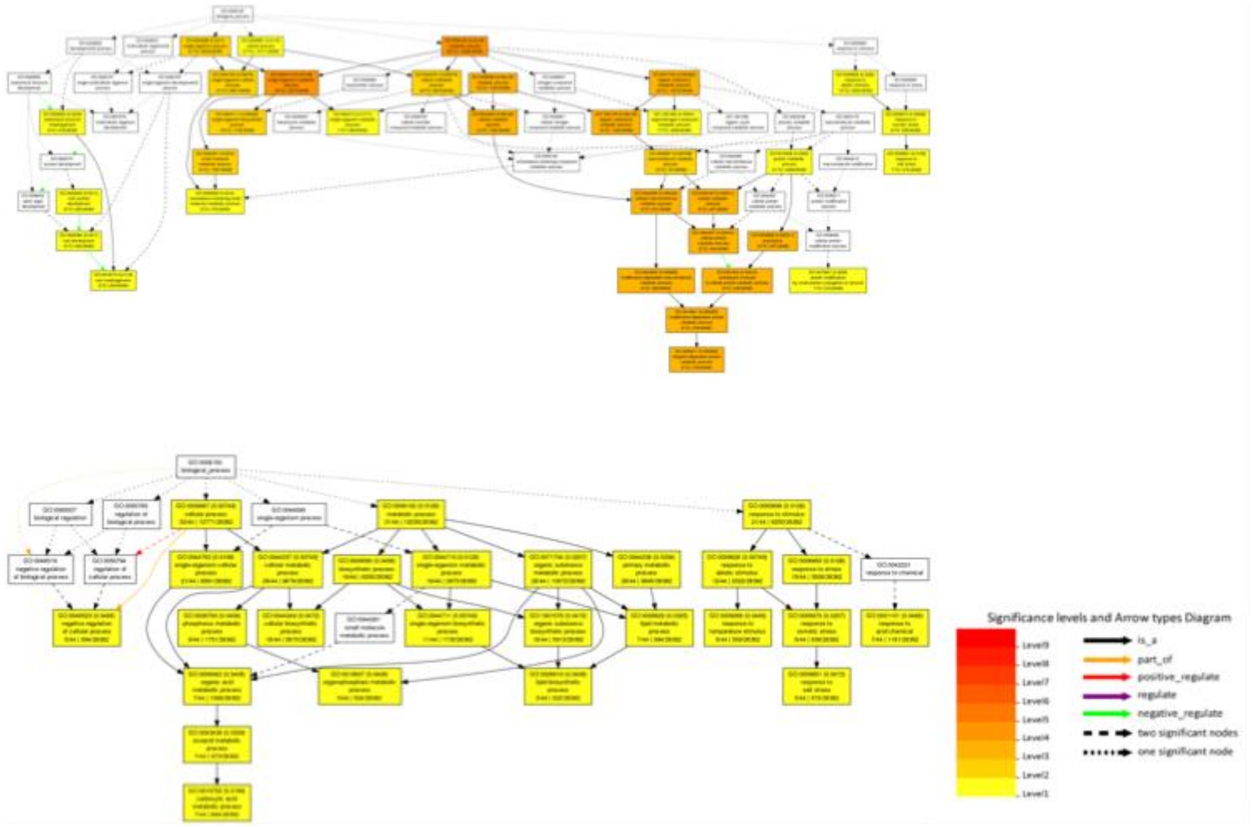


**Supplementary Figure 9. PANTHER pathway enrichment analysis of upregulated (top) and downregulated (bottom) of significantly regulated proteins from global proteomics are shown.**
